# Functional profiling of COVID-19 respiratory tract microbiomes

**DOI:** 10.1101/2020.05.01.073171

**Authors:** Niina Haiminen, Filippo Utro, Ed Seabolt, Laxmi Parida

## Abstract

In response to the ongoing global pandemic, progress has been made in understanding the molecular-level host interactions of the new coronavirus SARS-CoV-2 responsible for COVID-19. However, when the virus enters the body it interacts not only with the host but also with the micro-organisms already inhabiting the host. Understanding the virus-host-microbiome interactions can yield additional insights into the biological processes perturbed by viral invasion. With this aim we carry out a functional analysis of previously published RNA sequencing data of bronchoalveolar lavage fluid from eight COVID-19 patients, twenty-five community-acquired pneumonia patients, and twenty healthy controls. The resulting microbiome functional profiles and their top differentiating features clearly separate the cohorts. By examining the functional features in connection with their associated metabolic pathways, differentially abundant pathways are indicated, compared to both the community-acquired pneumonia and healthy cohorts. From this analysis, distinguishing signatures in COVID-19 respiratory tract microbiomes are identified, including decreased lipid and glycan metabolism pathways, and increased carbohydrate metabolism pathways. Here we present a framework for comparative functional analysis of microbiomes, the results from which can lead to new hypotheses on the host-microbiome interactions in healthy versus afflicted cohorts. The findings from this analysis call for further research on microbial functions and host-microbiome interactions during SARS-CoV-2 infection.

## Introduction

An impressive number of scientific studies have rapidly been published on the genomics and molecular-level host interactions of the new coronavirus SARS-Cov-2^1^ of reported bat origin,^2^ responsible for the COVID-19 disease pandemic. Additionally, understanding changes in the microenvironment within the host can yield insights into related biological processes perturbed by the virus. An increase in opportunistic organisms and an enhanced capacity for nucleotide and amino acid biosynthesis and carbohydrate metabolism has been associated with increased SARS-CoV-2 presence in fecal microbiomes.^3^ It has been suggested that gut microbiota could contribute to the pathology of the gut-lung axis during COVID-19.^4^ The respiratory microbiome members during the respiratory virus SARS-CoV-2 infection have also been studied.^1, 5–7^ Previous studies on the respiratory tract microbiome during other pathogen infections have examined its predictivity of clinical outcomes, and associated potential probiotic interventions.^8–12^

To better understand the role of the respiratory microbiome in COVID-19, we introduce a *functional* analysis, as opposed to taxonomic naming, of recently published metatranscriptomes^5^ from bronchoalveolar lavage fluid (BALF) of COVID-19 patients, healthy subjects, and community-acquired pneumonia (CAP) cases. While Shen et al.^5^ focused on the SARS-Cov-2 genomes and taxonomic profiling of the microbiomes, here we perform global functional profiling to characterize altered biological processes the respiratory tract microbiomes. Our protein domain focused amino acid matching approach supports the profiling of functions performed by known and potentially unknown organisms yet to be characterized.^13^ The robust comparative analysis presented here was designed to highlight consistent differences in COVID-19 patient microbiomes compared to both community-acquired pneumonia and healthy control samples.

## Results

### Functional profiling framework

The overall analysis workflow is depicted in Fig. 1A. The total RNA sequencing reads were first trimmed and filtered, followed by translation and functional classification with PRROMenade^14^ against a vast amino acid sequence collection of 21 million bacterial and viral protein domains from the IBM Functional Genomics Platform,^15^ annotated with KEGG enzyme codes (EC) from a corresponding functional hierarchy^16^ (see Supplementary Fig. S1 for filtering results). Post-processing and robust rank-based RoDEO^17^ projection onto a unified scale was performed to make the resulting functional profiles comparable.

**Figure 1.**
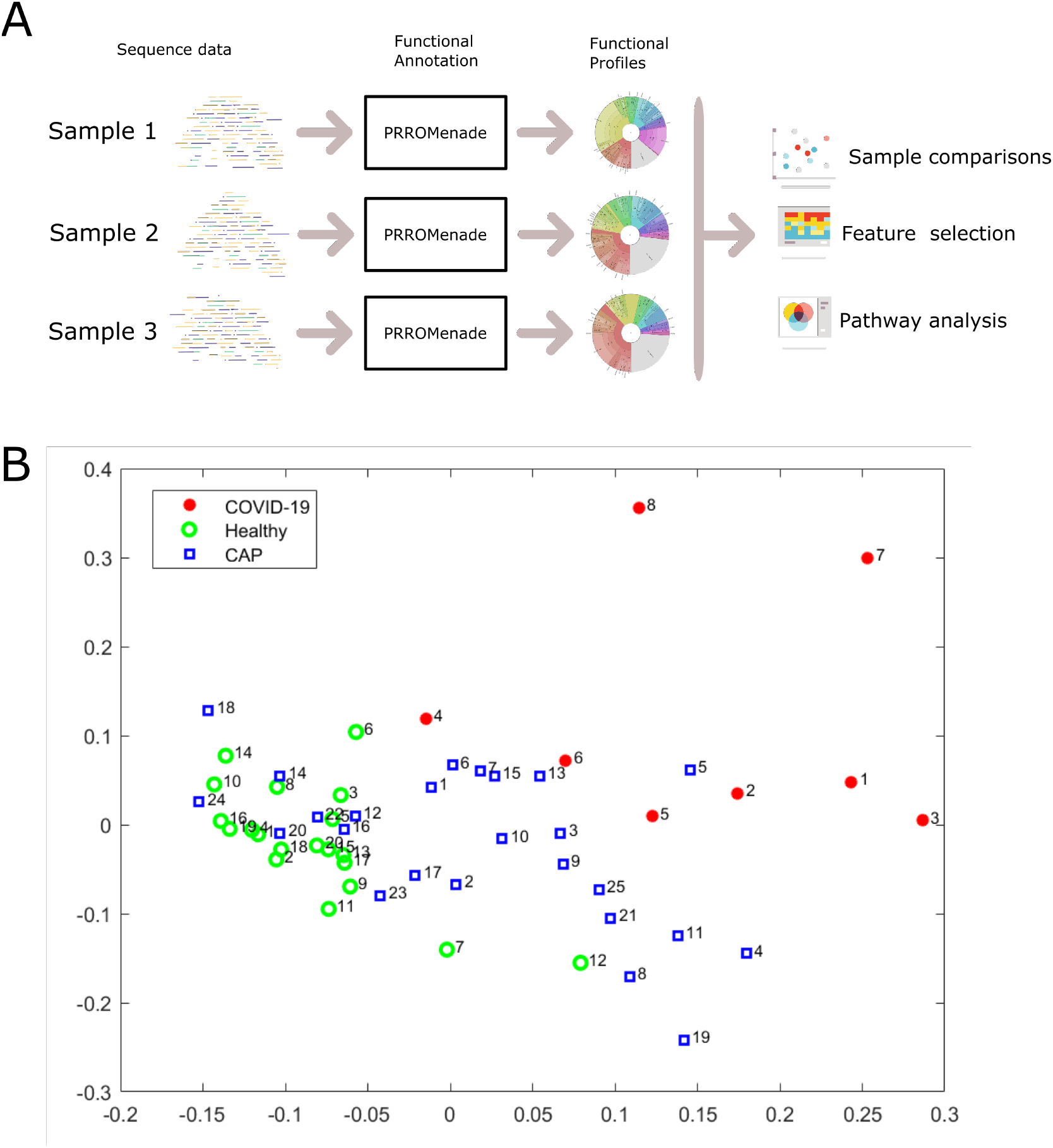
Overall analysis workflow and two-dimensional projection of functional profiles. **A**. Each microbiome sequencing sample is annotated with PRROMenade, utilizing labeled reference data from the IBM Functional Genomic Platform. The resulting functional profiles are visualized and compared in downstream analyses. **B**. Multidimensional scaling of the functional profiles using the Spearman distance. Each sample is represented by a marker colored by cohort and labeled by the sample number within that cohort.

### Microbiome functional profiles cluster by cohort

While the individual functional profiles vary, a robust comparative analysis reveals specific functions that are consistently altered between cohorts (Supplementary File S1 shows a Krona^18^ visualization of each sample). The read counts assigned at various functional hierarchy levels (Supplementary Fig. S2) were pushed down to the leaf level, and very low abundant features were removed for subsequent analyses (see Methods).

Multi-dimensional scaling of pairwise Spearman distances between the samples separates the COVID-19 cohort, while CAP samples are located between healthy and COVID-19 samples (Fig. 1B). A significant difference between the functional profiles was observed between the COVID-19, CAP, and healthy control cohorts according to the PERMANOVA test (*p* ≤ 0.0001). Functional profiling separated the cohorts with a similar score as taxonomic profiling (functional profiling *R*^2^ = 0.06 vs. taxonomic profiling by Shen et al^5^ *R*^2^ = 0.07).

### Differentially abundant features distinguish COVID-19 samples

The RNA sequencing data had varying total number of reads and human content per sample (Supplementary Fig. S1). Therefore we used RoDEO^17^ to project the functional profiles onto a robust uniform scale. To examine the most differentiating features for COVID-19 versus the other cohorts, we extracted 30 top-ranked features from the COVID vs. CAP comparison and from the COVID vs. healthy controls comparison. We then considered the union of the feature sets, resulting in 44 EC features.

When clustering the samples using the top differentiating features, the COVID-19 samples are grouped together and separate from the other cohorts, except for sample 4 (Fig. 2). While the examined 44 features were selected as those differentiating COVID-19 from CAP and healthy controls, they also separate the healthy control samples from all others; the healthy controls cluster tightly together. The results also demonstrate that the samples do not merely cluster by the total number of input reads or the fraction of functionally annotated microbial reads, since those measures vary within cohorts (Supplementary Fig. S1). The CAP patient samples were collected from different hospital sources, prior to the current pandemic, and represent pneumonia cases with various viruses detected in the sequencing data^5^ (e.g. enterovirus, influenza virus, rhinovirus), possibly contributing to the greater variability between their microbiome functional profiles.

**Figure 2.**
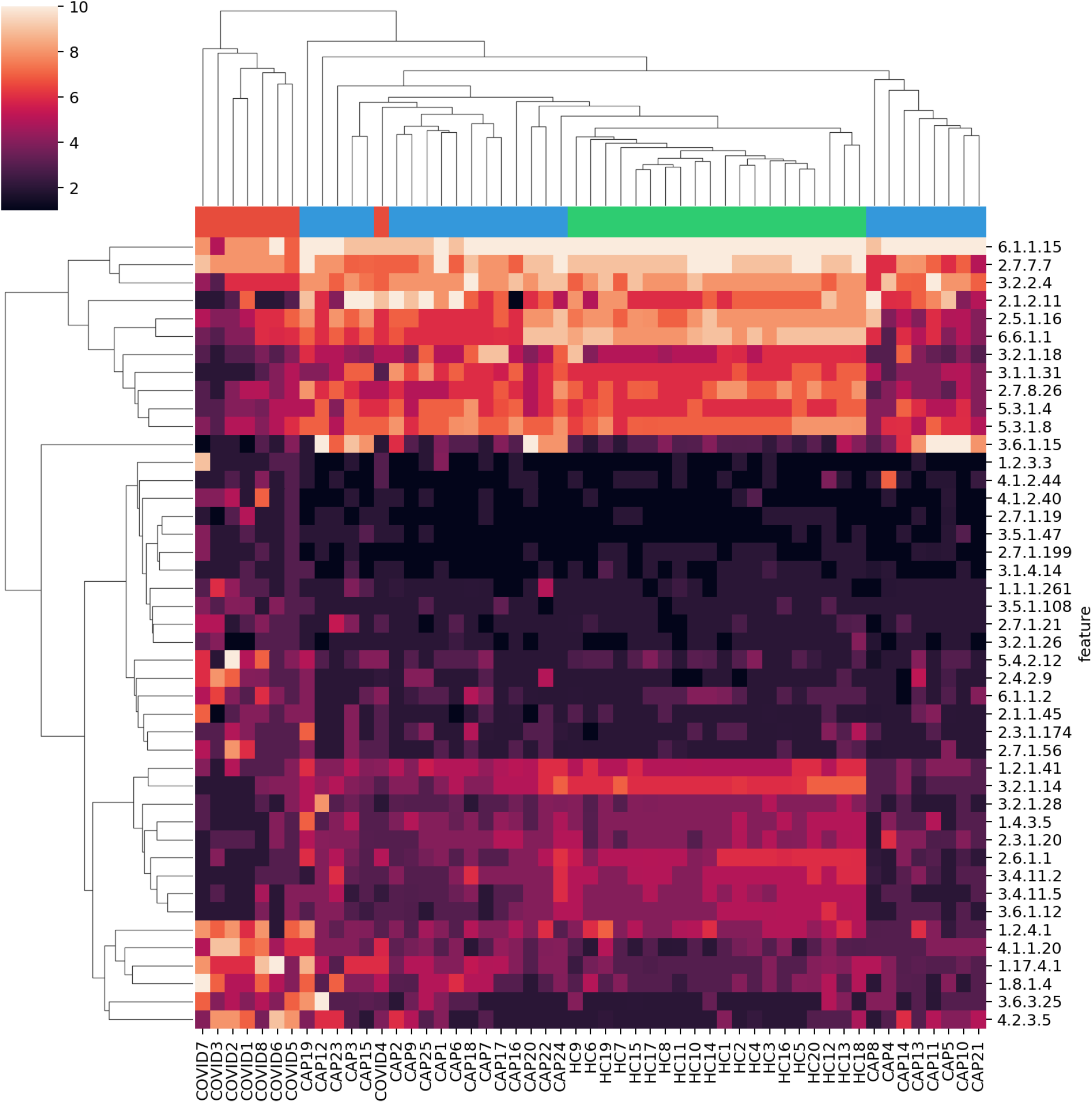
Clustering with top differentiating functional features. RoDEO processed EC abundance values (10 denotes highest possible value), for 44 features differentiating COVID-19 from community-acquired pneumonia and healthy controls. Columns and rows are ordered independently by hierarchical clustering of features and samples. The colors attached to the dendrogram on top reflect the cohort labels: red = COVID-19, blue = CAP, green = healthy control.

The COVID-19 samples have more abundant EC features including (see bottom left feature cluster in Fig. 2) 1.2.4.1 “Pyruvate dehydrogenase”, 4.1.1.20 “Diaminopimelate decarboxylase”, 1.17.4.1 “Ribonucleoside-diphosphate reductase”, 1.8.1.4 “Dihydrolipoyl dehydrogenase”, 3.6.3.25 “Sulfate-transporting ATPase”, and 4.2.3.5 “Chorismate synthase”, linked to various amino acid, carbohydrate, energy, and nucleotide metabolism pathways. EC 4.1.1.20 was also detected as increased in a metaproteome study of COVID-19 respiratory microbiomes.^19^ Supplementary Fig. S3 includes a scatter plot of the average change per EC in COVID-19 compared to CAP and healthy cohorts, highlighting outliers.

### Altered lung microbiome pathways indicated in COVID-19

In order to systematically examine the detected features against functional pathways, all the EC features were considered against their corresponding pathways from the KEGG metabolic pathway mapping.^16^ Pathway scores (mean abundance change in COVID-19) were computed using all the detected EC features per pathway, see Fig. 3. To identify outlying pathway scores (high or low compared to the observed distribution), median absolute deviation (MAD),^20^ a robust measure of dispersion was utilized. The most differential pathways are shown in Table 1.

**Figure 3.**
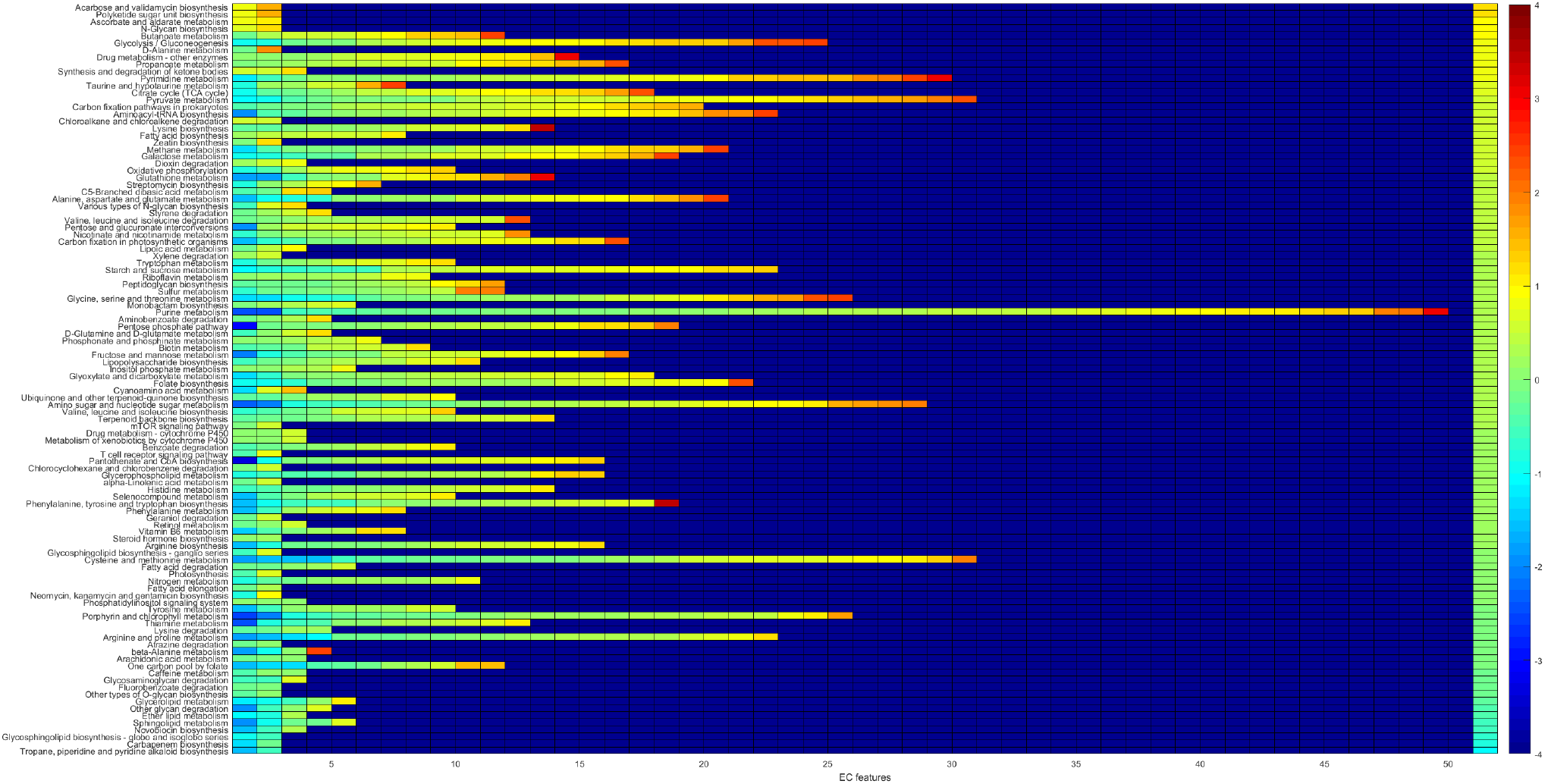
Pathway changes in COVID-19 samples. For each pathway (row), there are as many entries as there are detected EC features. The color of the entries indicate the average of COVID-19 vs. CAP and COVID-19 vs. HC changes. The entries on each row are ordered from low to high values. The background value (dark blue) indicates no data; some pathways have more detected features than others (only pathways with at least two EC features detected are considered). The rightmost column indicates the pathway score, the pathways are ordered accordingly from top (higher in COVID-19) to bottom (lower in COVID-19).

**Figure 4.**
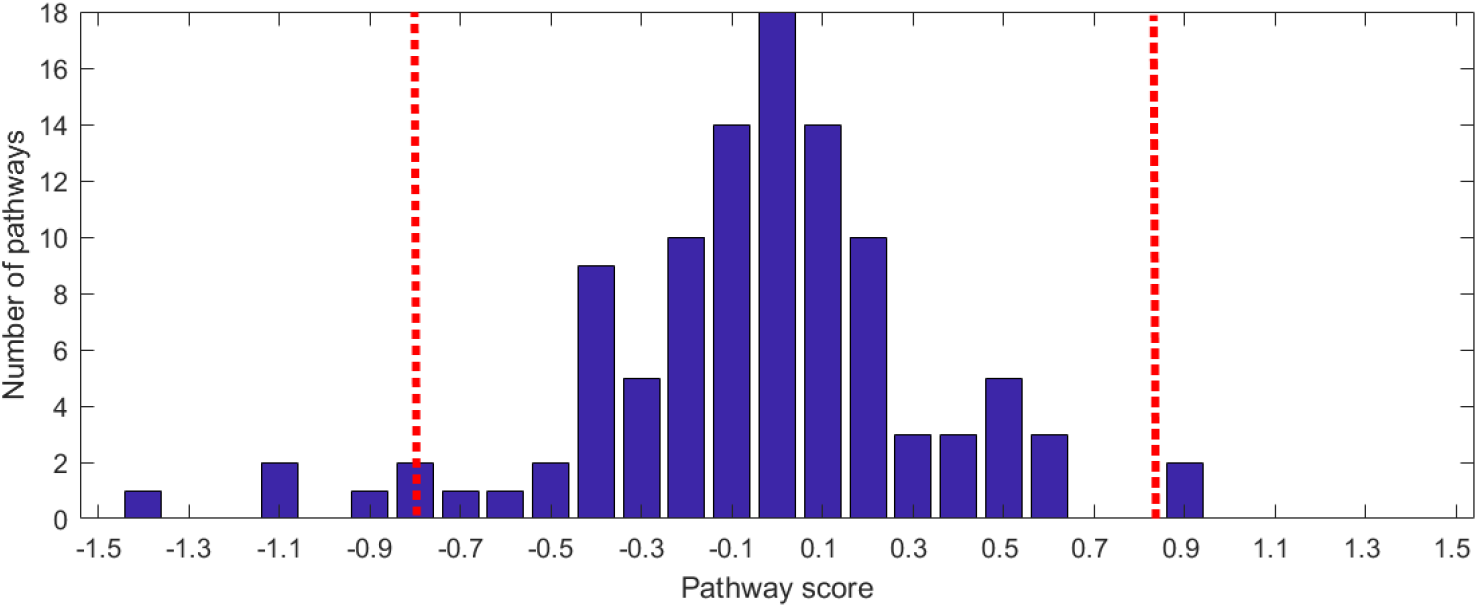
Pathway score distribution. The histogram of observed pathway scores is shown. The score thresholds for determining outliers according to median absolute deviation (MAD) is also marked (red dashed lines, three scaled median deviations away from the median).

**Table 1.** Altered pathways in COVID-19. The most differential pathways in COVID-19 (with dist. ≥2) are shown in the table. Here dist. denotes the pathway score’s distance from the median, divided by the scaled median absolute deviation. Pathways that are determined outliers (dist. ≥3) are marked with *. The direction of the change in COVID-19 (+/−), the number of associated EC features per pathway detected from functional profiling (#EC), and the pathway type are also shown.

| Pathway name | dist. | +/- | #EC | Type |
| --- | --- | --- | --- | --- |
| Tropane, piperidine and pyridine alkaloid biosynthesis | 5.04* | - | 2 | Biosynthesis of other secondary metabol. |
| Carbapenem biosynthesis | 3.84* | - | 2 | Biosynthesis of other secondary metabol. |
| Glycosphingolipid biosynth. - globo and isoglobo series | 3.75* | - | 2 | Glycan biosynthesis and metabolism |
| Novobiocin biosynthesis | 3.28* | - | 3 | Biosynthesis of other secondary metabol. |
| Sphingolipid metabolism | 2.90 | - | 5 | Lipid metabolism |
| Ether lipid metabolism | 2.82 | - | 3 | Lipid metabolism |
| Other glycan degradation | 2.64 | - | 4 | Glycan biosynthesis and metabolism |
| Glycerolipid metabolism | 2.11 | - | 5 | Lipid metabolism |
| Acarbose and validamycin biosynthesis | 3.47* | + | 2 | Biosynthesis of other secondary metabol. |
| Polyketide sugar unit biosynthesis | 3.47* | + | 2 | Metabolism of terpenoids and polyketides |
| Ascorbate and aldarate metabolism | 2.45 | + | 2 | Carbohydrate metabolism |
| N-Glycan biosynthesis | 2.38 | + | 2 | Glycan biosynthesis and metabolism |
| Butanoate metabolism | 2.33 | + | 11 | Carbohydrate metabolism |
| Glycolysis / Gluconeogenesis | 2.09 | + | 24 | Carbohydrate metabolism |
| D-Alanine metabolism | 2.04 | + | 2 | Metabolism of other amino acids |

Among the pathways that were lower in COVID-19 (Table 1) several are related to glycan biosynthesis and metabolism (e.g. other glycan degradation) and lipid metabolism (e.g. sphingolipid metabolism). Sphingolipids are important components of biomembranes, mediating signal transduction and immune activation processes, and they have been shown to decrease in COVID-19 patient sera.^21^ The feature 3.2.1.22 alpha-galactosidase (alpha-gal) is lower in COVID-19 and is linked to several of the lower abundance pathways in Table 1: glycosphingolipid biosynth. - globo and isoglobo series, sphingolipid metabolism, and glycerolipid metabolism. It has recently been hypothesized that dysbacteriosis observed in COVID-19 patients is linked to the reduction in the microbiota of alpha-gal containing commensal bacteria, or alternatively individuals with higher alpha-gal content in the microbiota may be less susceptible to COVID-19, supported by detected negative correlation between anti-alpha-gal antibody titers and COVID-19 disease severity.^22^ Elsewhere, raising anti-alpha-gal titers in the population by immunizing against inactivated harmless bacteria that harbor alpha-gal epitopes has been suggested.^23^ Here we additionally identify glycosaminoglycan degradation as decreased in COVID-19 samples (Fig. 3), while a connection between decreased presence of host glycosaminoglycan heparan sulfate modifying bacteria and increased COVID-19 susceptibility has been suggested.^24^

Among the pathways higher in COVID-19 are several related to carbohydrate metabolism, e.g. glycolysis/gluconeogenesis (Table 1). Enhanced microbial capacity for carbohydrate metabolism (glycolysis II from fructose-6-phosphate) has previously been indicated in fecal samples with a signature of high SARS-Cov-2 infectivity, along with decreased abundance of short-chain fatty acid producing bacteria.^3^

## Discussion

Analyzing COVID-19 respiratory tract metatranscriptomes from a functional perspective offers an additional view into their differences from healthy controls and from subjects with community-acquired pneumonia (CAP). The comparative approach taken here aims to mitigate possible experimental variation and resulting biases within individual samples, by focusing on detecting robust and consistent differences in the detected microbial functions. The framework presented here includes read filtering, functional annotation with a protein domain database and enzyme hierachy, feature abundance projection to a comparable scale, and finally metabolic pathway scoring to indicate differentially abundant functionalities in COVID-19 microbiomes compared to healthy and CAP samples. As a result, we identified both enzyme code features and metabolic pathways that differentiate COVID-19 respiratory tract microbiomes. The resulting functional profiles distinguish COVID-19 samples from others, similarly to the original taxonomy-based analysis of the community members.^5^

The differentially abundant functions and pathways identified here match findings from previous re ports, relating to decreased lipid and glycan metabolism,^21^ increased carbohydrate metabolism,^3^ and other characteristics of the microbiome linked to COVID-19.^22–24^ The remaining findings can suggest novel host-microbiome interactions to be examined further. Limitations of the study include small sample size and lack of clinical data, hence possible connections between administered therapies and the microbiome remain elusive. The findings from this analysis call for further in-depth research on microbial functions and host-microbiome interactions during SARS-CoV-2 infection. Examining metatranscriptome sequencing reads with this comparative functional annotation framework could yield additional insights into microbiome alterations also in other diseases.

## Methods

### Sequence data and functional database

The recently published bronchoalveolar lavage fluid (BALF) metatranscriptomic sequencing data^5^ of 8 COVID-19 patients, 20 healthy controls (HC), and 25 cases of community-acquired pneumonia (CAP) were obtained from the National Genomics Data Center (accession PRJCA002202).^25^ Pre-processing included TrimGalore^26^ adapter and quality trimming (–length 50 ‒trim-n–max_n 10), and poly-A trimming performed with BBduk^27^ (trimpolya=10, minlength=50). The reads were filtered against human (GCF_000001405.39), the PhiX sequencing control (GCF_000819615.1), and the SARS-CoV-2 virus (NC_045512.2) with BBsplit^27^ (ambiguous=random, ambiguous2=split). The paired reads were processed separately, individual reads that did not match the human, PhiX, or SARS-CoV-2 genomes (278k to 50.7M reads per sample) were retained for the microbial community functional annotation (see Supplementary Fig. S1 for the filtering results).

The KEGG Enzyme Nomenclature (EC) reference hierarchy^16^ was used as the functional annotation tree. The EC numbers define a four-level hierarchy. For example, 1.5.1.3. = “Dihydrofolate reductase” is a fourth (leaf) level code linked to top level code 1 = “Oxidoreductases”, via 1.5. = “Acting on the CH-NH group of donors” and 1.5.1 = “With NAD+ or NADP+ as acceptor”. A PRROMenade^14^ database search index was constructed using the KEGG hierarchy and a total of 21.2M bacterial and 53k viral annotated protein domain sequences (of minimum length 5 AA), obtained on June 6, 2020 from the IBM Functional Genomics Platform^15^ (previously known as OMXWare). An earlier release of the bacterial domain data has been discussed previously in conjunction with PRROMenade indexing.^14^

### Functional annotation and analysis

Metatranscriptomic sequencing reads were annotated with PRROMenade by locating the maximal length exact match for each read, and processed as described previously.^14^ Minimum match length cutoff of 11 AA (corresponding to 33 nt) was employed. Classified (non-root) read counts (6.8k to 11.5M per sample, see Supplementary Fig. S1–S2) were post-processed to summarize the counts at the leaf level of the functional hierarchy. Leaf nodes contributing ≥ 0.05% of total annotated reads in at least one sample were retained, resulting in 633 leaf nodes to include as the features of the functional profiles. Multidimensional scaling (Matlab function cmdscale, *p* = 2) and permutational multivariate analysis of variance (f_permanova, *iter* = 100, 000, from the Fathom toolbox^28^ for Matlab) were applied on pairwise Spearman’s distances (Fig. 1B).

Subsequently, the profiles were processed with RoDEO^17^ (*P* = 10, *I* = 100, *R* = 10^7^) for robust comparability. The per-sample parameter *P′* was determined according to the number of annotated reads as previously described (in Supplementary File 2 by Klaas et al.^29^). A two-sample Kolmogorov-Smirnov test (kstest2 in Matlab) was applied to identify differentially abundant features between COVID-19 samples and CAP, healthy control samples. Features were ordered by p-value and top features selected for average linkage hierarchical clustering using the Euclidean distance (Fig. 2).

### Pathway analysis

The KEGG^16^ metabolic pathway maps were utilized to link functions with pathways, and the pathways were analyzed for changes between COVID-19 and CAP, HC. The pathways were evaluated for average abundance change as follows. Let *a*_*i*_ be the mean RoDEO abundance of EC feature *i* for COVID-19 samples, *b*_*i*_ for CAP samples, and *c*_*i*_ for HC samples. The feature score is defined as *fs*_*i*_ = ((*a*_*i*_ − *b*_*i*_) + (*a*_*i*_ − *c*_*i*_))/2, positive values indicating higher abundance in COVID-19. Pathway score 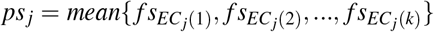 was computed using the set of features, *EC*_*j*_, that map to the pathway *j* (considering only pathways with *k* ≥ 2). The pathway score distribution was normalized to have mean zero for visualization.

Median absolute deviation (MAD)^20^, a robust measure of dispersion, was used to identify outliers from the observed pathway score distribution (isoutlier in Matlab with the parameter median). With the default parameters, an outlier is defined as a value that is more than three scaled median absolute deviations away from the median.

## Author contributions

N.H. and F.U. conducted the experiments, analyzed the results and wrote the manuscript. E.S. provided reference data from the IBM Functional Genomics Platform. N. H. and L.P. conceived the study and analyzed the results. All authors reviewed the manuscript.

