## Supplemental Figures for "Functional profiling of COVID-19 respiratory tract microbiomes"

### Supplementary Figures S1—S3

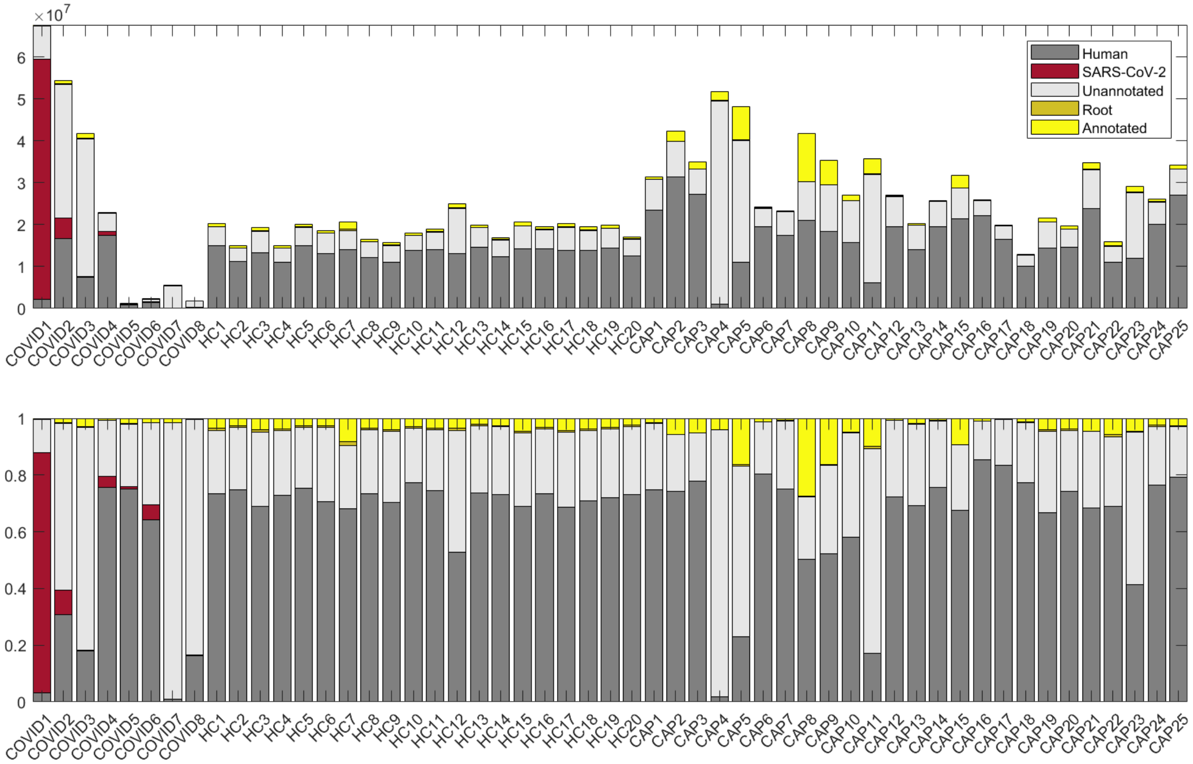


**Supplementary Figure S1:** Distribution of the read counts (after quality- and PhiX-filtering) for COVID-19 (COVID), Healthy Control (HC) and community acquired pneumonia (CAP) samples. Annotated reads denote those with a match in the protein domain database below the root level, reads assigned to the root are counted separately (see also Figure S2 for the annotated read levels in the functional hierarchy). **Top**: The number of reads per sample, including human- and SARS-CoV-2 matching reads. Each read in a pair was evaluated and counted separately. **Bottom**: Same data as above, scaled to indicate fractions from the total read count per sample.

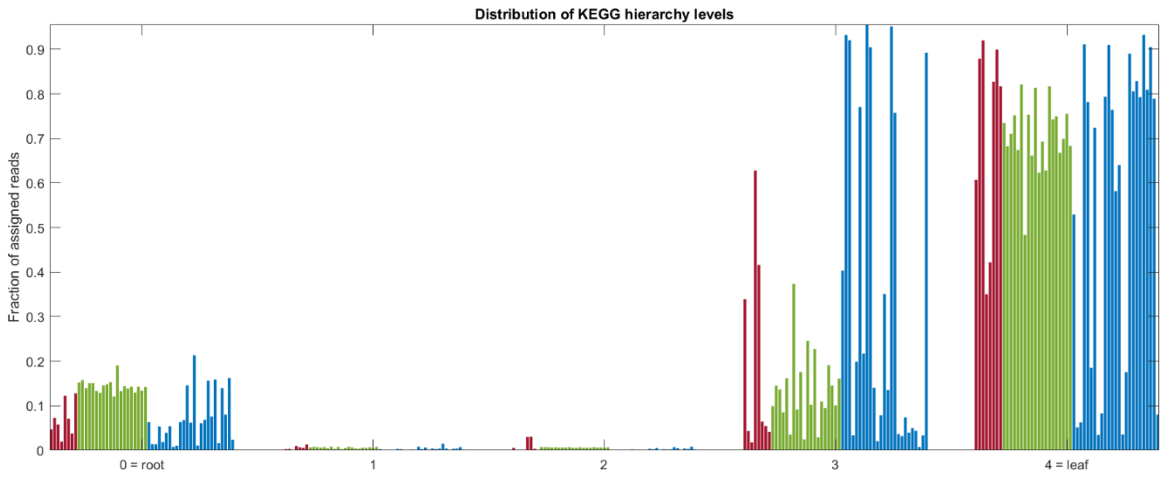


**Supplementary Figure S2.** Fraction of (human-filtered & SARS-CoV-2 filtered) reads assigned to each functional hierarchy level, one bar per sample: COVID-19 (red), healthy control (green), community-acquired pneumonia (blue).

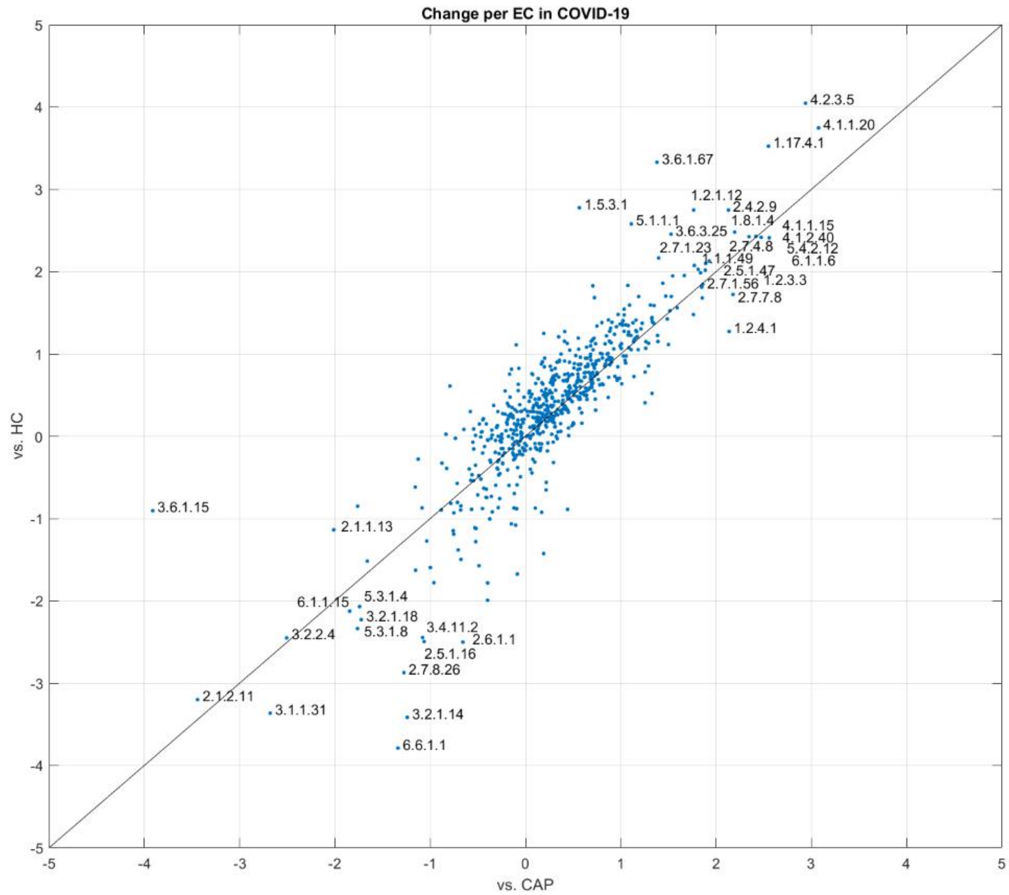


**Supplementary Figure S3.** The mean RoDEO abundance change for each EC (dot in the figure) in COVID-19 vs. CAP (x-axis) and vs. Healthy controls (y-axis). The features with a change of at least 2 in either comparison are labeled.
